# The microbiota plays a critical role in the reactivity of lung immune components to innate ligands

**DOI:** 10.1101/2020.10.19.345116

**Authors:** Quentin Marquant, Daphné Laubreton, Carole Drajac, Elliot Mathieu, Edwige Bouguyon, Marie-Louise Noordine, Aude Remot, Sabine Riffault, Muriel Thomas, Delphyne Descamps

**Author notes:** Co-authors (Quentin Marquant, Daphné Laubreton). Corresponding author, **Correspondence to:** Delphyne Descamps. Virologie et Immunologie Moléculaires, INRAE, UVSQ, Domaine de Vilvert, 78350 Jouy-en-Josas, France.

## Abstract

The microbiota contributes to shaping efficient and safe immune defenses in the gut. However, little is known about the role of the microbiota in the education of pulmonary innate immune responses. Here, we tested whether the endogenous microbiota can modulate reactivity of pulmonary tissue to pathogen stimuli by comparing the response of specific pathogen-free (SPF) and germ-free (GF) mice. Using SPF and GF mice intranasally exposed to lipopolysaccharide (LPS), a component of Gram-negative bacteria, we observed earlier and greater inflammation in the pulmonary compartment of GF mice than that of SPF mice. Toll-like receptor 4 (TLR4) was more abundantly expressed in the lungs of GF mice than those of SPF mice at steady state, which could predispose the innate immunity of GF mice to strongly react to environmental stimuli. Lung explants were stimulated with different TLR agonists or infected with the human airways pathogen, respiratory syncytial virus (RSV), resulting in greater inflammation under almost all conditions for the GF explants. Finally, alveolar macrophages (AM) from GF mice presented a higher innate immune response upon RSV infection than those of SPF mice. Overall, these data suggest that the presence of microbiota in SPF mice induced a process of innate immune tolerance in the lungs by a mechanism which remains to be elucidated. Our study represents a step forward to establishing the link between the microbiota and the immune reactivity of the lungs.

**Plain Language summary:** Microbiota represents an important partner of immunologic system at the interface between immune cells and epithelium. It is well known, notably in the gut, that the microbiota contributes in shaping efficient and safe defenses. However, little is known about the role of the microbiota in the education of pulmonary innate immune responses. In this study, we postulate that endogenous microbiota could dampen an excessive reactivity of pulmonary tissue to external stimuli. Thus, we sought to study the innate immune reaction switched on by viral or bacterial ligands in respiratory tract cells coming from mice with or without microbiota (germ-free condition, GF). Altogether, our results show a higher inflammatory reaction in GF condition. This study represents a step forward to better establish the link between the microbiota and the reactivity of the lung tissue. Not only these data demonstrate that the microbiota educates the pulmonary innate immune system, but also contributes the emerging concept of using respiratory commensal bacteria as potential next-generation probiotics to prevent susceptibility to respiratory diseases.

## Introduction

The holobiont defines the host and its associated communities of microorganisms in tissues such as the gut, skin, vagina, and lungs. An integrated view of holobiont assembly, stability, and function provides a clue to the better control of symbiosis and the prevention of dysbiosis (1). Aside from global approaches to study systemic functional mechanisms in a whole organism, an understanding of the holobiont can also be gained from studies of specific tissues, as exemplified by studies of the gut. Numerous studies showing that the arrival of the microbiota affects immunity and shapes the epithelium in the gut have been conducted (2). In this field, the use of GF animals has aided determination of the genuine morphology, immunity, and maturation of tissues, before any contact with microorganisms (3).

It is now widely accepted that bacteria colonize the airway compartment at birth (4), suggesting that the lungs may be functionally influenced by the microbiota. In mice, culture methods have shown the lung microbiota to represent an estimated 10^3^-10^5^ CFU/g of lung tissue (5). Dickson *et al*. proposed that the lower airway microbiota is the result of bacterial migration through a transition process of oropharyngeal secretions, micro-aspiration, and direct dispersal of mucosa from the upper respiratory tract to the lower compartment (6–8). The airway microbiota is controlled by a balance between growth (if any), immigration, and elimination processes (9). How the microbiota shapes pulmonary innate immune responses is poorly understood, despite a number of pioneering studies. According to Herbst *et al*., the GF status exacerbates the allergic responsiveness triggered by intraperitoneal treatment with ovalbumin of adult GF mice. Conversely, the colonization of previously GF mice reduced their allergic response (10). Dysbiosis of the upper airway microbiota has been reported during the course of chronic respiratory diseases, such as asthma (11, 12), and also upon respiratory viral infections, such as RSV that causes bronchiolitis in infants (13, 14). In addition, development of the microbiota, and consequently exposure to bacterial components in early life represents an important step in the immune history of individuals. Indeed, children living in rural areas are exposed to a larger diversity of microbes and/or to environments rich in LPS, which protects them from the development of asthma (15, 16), or from an excessive inflammatory response to RSV infection (17). Interestingly, *TLR4* and *CD14*, two co-receptors of LPS, and *TLR2* mRNA expressions are downregulated in the respiratory secretions of healthy Argentinian infants regularly exposed to LPS in their environment compared with infants from urban areas with low LPS levels (17). This supports the concept of a “window of opportunity”, consisting of a period in early life when the immune system can be shaped, notably by the microbiota, which influences the sensitivity of individuals to develop various diseases (18). Also, the administration of commensal respiratory bacteria (*Corynebacterium pseudodiphtheriticum* and *Dolosigranulum pigrum*) to the airways of infant mice protects them against pathogens in a strain dependent manner, such as RSV, or a secondary infection with *Streptococcus pneumoniae* (19, 20). Furthermore, the lungs are constantly challenged by microorganism signals, originating not only from the local microbial populations but also from a distance by signals sent from the gut microbiota through the gut-lung axis (21). Regardless of the source of the microbial signals, we postulate that the mechanisms of the lung innate immune responses and their intensity are conditioned by such microorganisms.

By comparing GF mice with their SPF counterparts, Gollwitzer *et al*. observed no significant differences in the frequency of dendritic cells (DCs) (13) (CD11b^+^ or CD103^+^) in the lungs and the expression of costimulatory molecules, such as CD40, ICOSL, CD80, and PD-L2 was not influenced by the presence or absence of the microbiota (22). Only the peak of PD-L1 expression on CD11b^+^ DCs was abrogated in GF mice during the first two weeks of life in the lungs but not on AMs (22). Our results have also shown there were no marked differences in the lung physiology or number of immune cells, although there were some differences in mRNA levels for certain innate genes between GF and SPF mice (5). All DCs, including plasmacytoid DCs (pDCs) and the two subsets of conventional DCs (13), CD4^+^ and CD8^+^ T cells, and B cells, are present in similar numbers in GF and SPF lungs. The global structure of the epithelium and main immune-cell subsets in the lungs are comparable between GF and SPF mice (23). The main actors of immunity appear to be well installed in the lungs of GF mice, but their function and activity have not been studied thus far, particularly in the presence of microbial stimuli that can mimic the first contact of the lungs with the environment at birth. Thus, we sought to compare the innate immune reaction of GF versus SPF mice, either *in vivo* or using freshly isolated pulmonary cells (lung explants and AMs) upon exposure to TLR agonists or to a respiratory pathogen, namely RSV. Overall, we observed a stronger inflammatory reaction in the lungs of GF mice, and in GF pulmonary explants and AMs. Such hypersensitivity of the innate immune system of GF lungs was also characterized by stronger anti-viral responses in infected AMs. Thus, we postulate that the microbiota matures lung immunity towards delayed and balanced responsiveness, potentially avoiding exacerbated innate immune responses.

## Materials and methods

### Mice

Male SPF C57Bl/6Jrj mice (8 weeks old) were purchased from Janvier (Le Genest, St. Isle, France) and bred in the Rodents and Fishes Experimental Infectiology Facility (IERP, INRAE Jouy-en-Josas, France). Male GF C57Bl/6Jrj mice (8 weeks old) were bred and housed under germ-free conditions in Trexler-type isolators (ANAXEM, INRAE, Jouy-en-Josas). All experiments were approved by the ethics committee COMETHEA (Ethical Committee for Animal Experimentation, INRAE, and AgroParisTech; authorization number: APAFIS#3441-2016010614307552vl).

### Virus

Human RSV-A2 was obtained from John Tregoning (Imperial College London, United kingdom). Recombinant human cherry-labelled RSV (RSV-Cherry) was produced as previously described (24).

### In vivo LPS administration

SPF and GF C57Bl/6Jrj mice were anesthetized with ketamine/xylazine (Rompun/Imalgene) and 10 ng/g LPS from *E.coli* O111: B4 (Sigma) or saline solution (PBS) were injected by the intranasal (i.n.) route. Mice were euthanized with pentobarbital at 1h30, 3h, and 6h post-treatment. Bronchoalveolar lavages (BAL) and lung tissues were collected for further analyses.

### Bronchoalveolar lavage

Cells were collected from BAL and centrifugated at 400 x g for 10 min at 4°C. May-Grunwald-Giemsa (MGG) coloration was performed to assess the differential cell counts using light microscopy. Two hundred cells were counted for determination of the relative percentage of neutrophils. Cell-free BAL samples were stored at −80°C for cytokine analysis.

### Ex vivo lung explant stimulation

Lung explants (150 μm) were obtained from fresh lungs of SPF and GF mice using a Krumdieck tissue slicer MD 6000 (Alabama Research and Development), as previously described (5). For TLR stimulation experiments, lung explants were stimulated with Poly(I:C) (100 μg/mL, Sigma-Aldrich), LPS (500 ng/mL, E.coli 011:B4, Sigma-Aldrich), Imiquimod (10 μg/mL, Invivogen), or CpG-B ODN1826 (10 μg/mL, Sigma) for 24 or 48h at 37°C. For viral infection experiments, lung explants from SPF or GF mice were infected with RSV-cherry (dilution = 1/100, corresponding to 5,480 pfu/well or 2.75 x 10^4^ pfu/mL) or Mock control for 2h, washed with complete RPMI medium, consisting of RPMI 1640 (Gibco) supplemented with 5% heat-inactivated fetal calf serum (Gibco), 2 mM L-glutamine (Gibco), and antibiotics (100 U/mL penicillin and 100 μg/mL streptomycin, Invitrogen), and incubated for 24 or 48h at 37°C. Supernatants were collected for cytokine measurements. Lung explants were lysed in 300 μL Lysis Buffer (5 mM EDTA, 150 mM NaCl, 50 mM Tris-HCl, Triton 1%, pH 7,4), containing anti-proteases (Roche), with a Precellys 24-bead grinder homogenizer (Bertin Technologies) for 2 × 15 s at 4,000 x g for protein extraction. Lung explant homogenates were clarified by centrifugation for 10 min at 10,000 x g and collected on microplates. Protein quantity in lung explant homogenates was measured using the DC™ Protein Assay (Bio-Rad), according to the manufacturer’s instructions.

### Ex vivo AMs culture and infection

BAL was performed on GF and SPF mice to collect AMs. Red blood cell lysis buffer (Sigma) was used to eliminate any blood cells. Cells were then centrifuged for 10 min at 400 x g at 4°C and counted on a microscope using Trypan-blue solution to exclude dead cells. Cells (1 x 10^5^) were plated on 96-well flat-bottom plates in RPMI complete medium, as previously described (25). After 1h, the medium was replaced to eliminate non-adhesive cells and debris. The purity of the AMs was routinely > 95%. After another 24h, AMs were infected with Mock control solution or human RSV-A2 (MOI = 5) in RPMI medium without serum. The medium was renewed 2h post-infection and replaced with RPMI complete medium. Supernatants and cells were collected 24h post-infection for analysis.

### Cytokine measurement

Cytokine production was measured by ELISA using antibodies against IL-5, IL-12p70, IFNg, IL-4, IL-6, IL-10, IL-22, IL-25, Tumor-necrosis factor *α* (TNFα), Thymic stromal lymphoprotein (TSLP) (all Ready-Set-Go!^®^ ELISA, ThermoFisher Scientific), and KC/CXCL1 (R&D Systems), according to manufacturers’ instructions. Concentrations were normalized to the quantity of protein in the lung explant homogenates or the weight of the lungs in lung homogenates. IFNα and IFNβ measurements were assessed in AMs supernatants using the IFNα/IFNβ 2-Plex Mouse ProcartaPlex™ immunoassay (ThermoFisher), according to the manufacturer’s instructions and read on a MagPix instrument (Luminex). Data were analyzed using Bio-Plex Manager software (Bio-Rad).

### Cytotoxicity

The cytotoxicity in lung explants was measured using the Non-Radioactive Cytotoxicity Assay kit (Promega), according to the manufacturer’s instruction, and determined as follows: cytotoxicity (%) = (OD490 of LDH in the supernatant)/(OD490 of LDH in the supernatant + OD490 of LDH in the explant lysate) x 100.

### RT-qPCR

Total RNA was extracted from lung lysates or AMs using a NucleoSpin^®^ RNA midi or XS column (Macherey Nagel), respectively, and reverse transcribed using random primers and reverse transcriptase (SuperScript II, Invitrogen), according to the manufacturer’s instructions. The primers (Sigma–Aldrich) used are listed in Table 1. RT-qPCR was performed in duplicate for each gene using the CFX Connect™ Real-Time PCR Detection System (Bio-rad) and SYBRGreen PCR Master Mix (Eurogenetec). Data were analyzed using CFX Manager™ Software (Bio-rad) to determine the quantification cycle (Cq) values. Results were determined using the formula 2^-ΔCq^, with ΔCQ = Cq_gene_-Cq_GADPH or HPRT_.

**Table 1.**
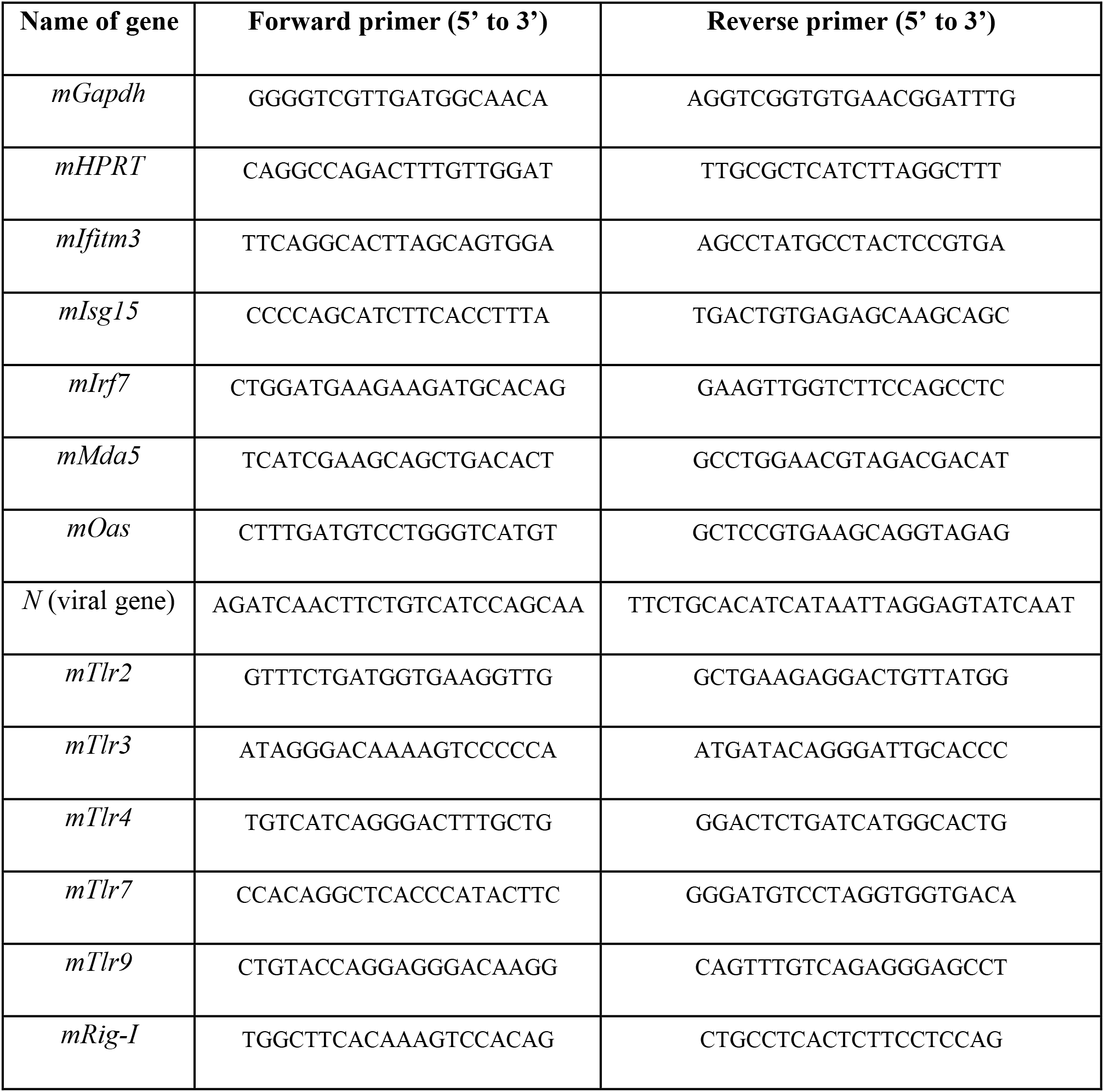
List of primers used for RT-qPCR.

### Western blots

The lungs of GF and SPF mice were crushed in Lysis Buffer (5 mM EDTA, 150 mM NaCl, 50 mM Tris-HCl, 1%, Triton pH 7,4) using a Tissue Homogenizer System device (Precellys 24). Lung lysates were then centrifugated at 10,000 x g and the supernatants collected. Lowry’s procedure (Bio-Rad) was used for protein assays. Western-blot analysis was performed by 12% SDS-PAGE using anti-mouse TLR4 (sc-30002, Santa Cruz), with an appropriate peroxidase-conjugated secondary antibody (Jackson ImmunoResearch Laboratories). Protein loading was determined using an anti-β-actin (AC-74, Sigma-Aldrich) antibody. Signals were detected on a ChemiDoc XRS+ device (Bio-Rad) and analyzed using Image Lab software (Bio-Rad).

### Data and statistical analysis

For each condition and cytokine, the ratios shown in the heatmaps were obtained using the following formula: ratio = (average of the GF stimulated conditions – average of the control conditions) / (average of the SPF stimulated conditions – average of the control conditions). Ratios were entered into Morpheus online software (Broad Institute) to obtain the heatmaps. Column statistics using the D’Agostino and Pearson test was performed to determine whether the data come from a Gaussian distribution before running the statistical analysis. Differences between the experimental groups were statistically evaluated using a non-parametric Mann-Whitney test (non-Gaussian distribution; groups = 2). All tests were performed using Graphpad Prism, Version 8 for Windows (GraphPad Software Inc). Data are expressed as the mean ± SD. Statistically significant differences were defined as *\*P* < 0.05, \*\**P* < 0.01, \*\*\**P* < 0.001, and \*\*\*\**P* < 0.0001.

## Results

### The lung inflammatory reaction triggered by a single LPS intranasal administration is enhanced in GF mice

We characterized the innate immune response of pulmonary tissue by intra-nasally (i.n.) treating SPF and GF mice with LPS, a major component of the outer membrane of Gram-negative bacteria, or PBS as control. Pro-inflammatory markers were measured at various time points (1h30, 3h, and 6h) post-instillation of LPS in the airways to follow the kinetics of the response (Fig. 1A). LPS treatment induced strong production of TNFα, IL-6, and KC/CXCL1, detectable in bronchoalveolar lavages (BAL) and lung homogenates, both in GF and SPF mice, relative to their basal levels after the administration of PBS. The production of TNFα was stronger in BAL and lung homogenates of GF mice than that of SPF mice at the earliest time point (1h30) (Fig. 1B). IL-6 production was higher in the BAL of GF mice at 1h30 and in the lung homogenates at 3h (Fig. 1C). Levels of the neutrophil attracting chemokine, keratinocyte chemoattractant (KC/CXCL1), were also higher in BAL and lung homogenates of GF mice at 1h30. The levels of KC/CXCL1 remained higher in the lungs of GF than SPF mice at 3h and 6h after LPS administration (Fig. 1D). This observation correlated with the earlier recruitment of neutrophils in the BAL of GF than that of SPF mice (3h *vs* 6h) (Fig. 1E). Overall, our data show that the lungs of GF mice exhibit more rapid kinetics (for all markers) and increased immune reactivity (for KC/CXCL1 and neutrophils in BAL) to LPS exposure.

**Figure 1.**
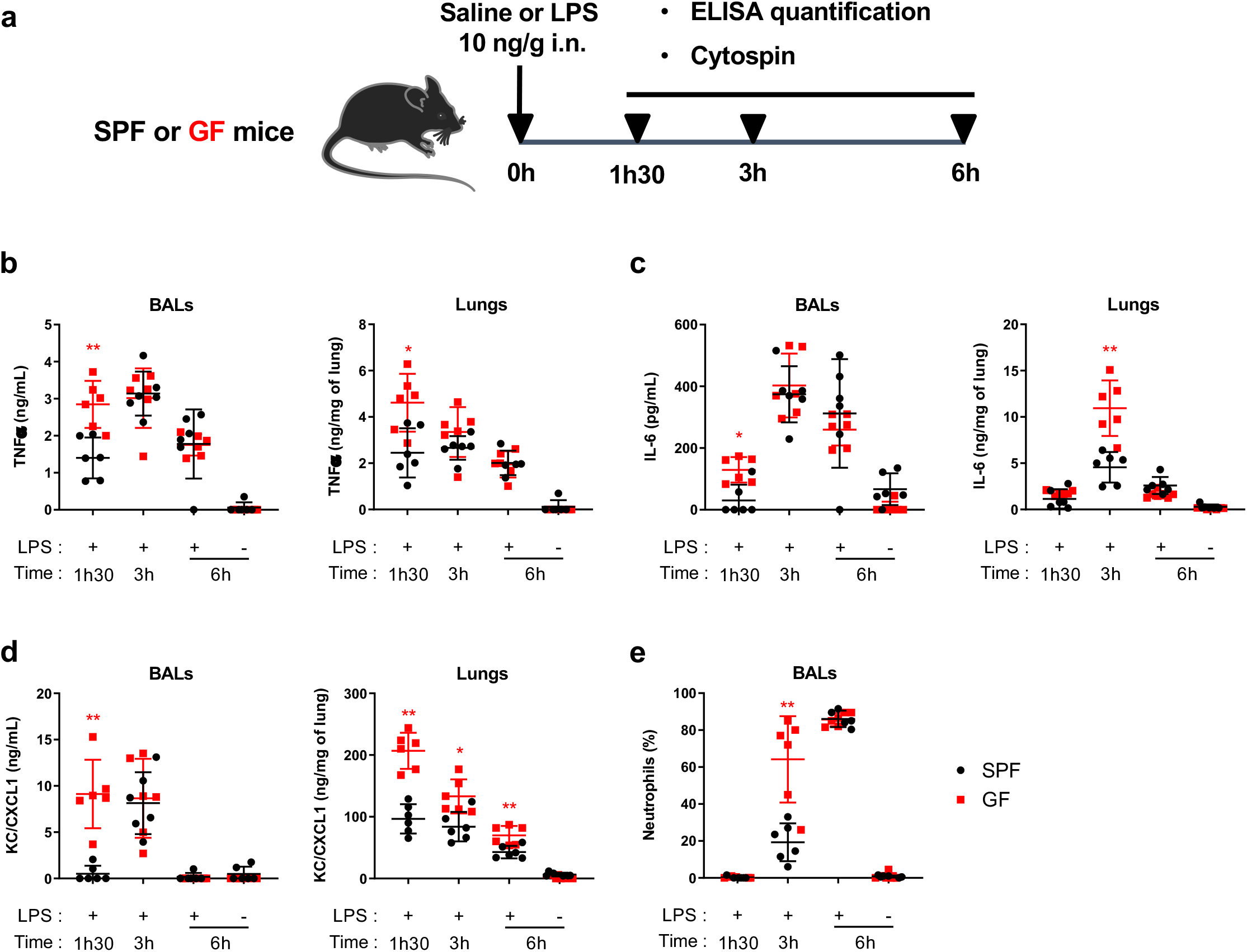
GF mice show earlier upregulation of pro-inflammatory markers in the lungs after LPS stimulation. (**A**) Experimental design. SPF or GF mice were anesthetized and saline solution or LPS (10 ng/g) administrated intra-nasally (i.n.). Mice were euthanized 1 h 30, 3 h, or 6 h after LPS administration to evaluate pulmonary pro-inflammatory parameters. (**B**) TNFα, (**C**) IL-6, and (**D**) KC/CXCL1 production was assessed in BAL (ng/mL) and the lung homogentaes (ng/mg of tissues) of SPF and GF mice. (**E**) The percentage of neutrophils was also determined in BAL. Results are expressed as the mean ± SD (n = 6 mice/group) and are from two independent experiments. \**P* ≤ 0.05, \*\**P* ≤ 0.01 between the SPF and GF groups (Mann-Whitney test) at the same time post-instillation (**B**-**E**).

### Steady-state TLR4 expression in the lungs is higher in the absence of microbiota

TLR4 is the key sensor that initiates the inflammatory response to LPS exposure (26). Indeed, mice deficient in TLR4 expression (TLR4^-/-^ mice) were unable to produce an inflammatory response following LPS instillation (Fig. S1). In a first step to understand why the lung tissue response to LPS was enhanced in the absence of microbiota, we analyzed TLR4 expression in the lungs of SPF and GF mice at steady state by western blotting (Fig. 2A). In these experimental conditions, TLR4 protein was detectable in the lungs of GF mice but not in the lungs of SPF mice. No TLR4 protein was detected in the lungs of TRL4^-/-^ mice, demonstrating the specificity of the antibody (Fig. 2B). In our hands, none of the other TLRs were detectable by western blotting (data not shown). The relative expression of mRNA levels of several pattern-recognition receptors (PRRs), namely *Tlr3, Tlr4, Tlr7, Tlr9, Rig-1*, and *Mda5* were similar in the lungs of both SFP and GF mice at steady state (Fig. 2C). These data suggest that the stronger pulmonary innate immune response to LPS in GF mice may be related to higher TLR4 protein expression.

**Figure 2.**
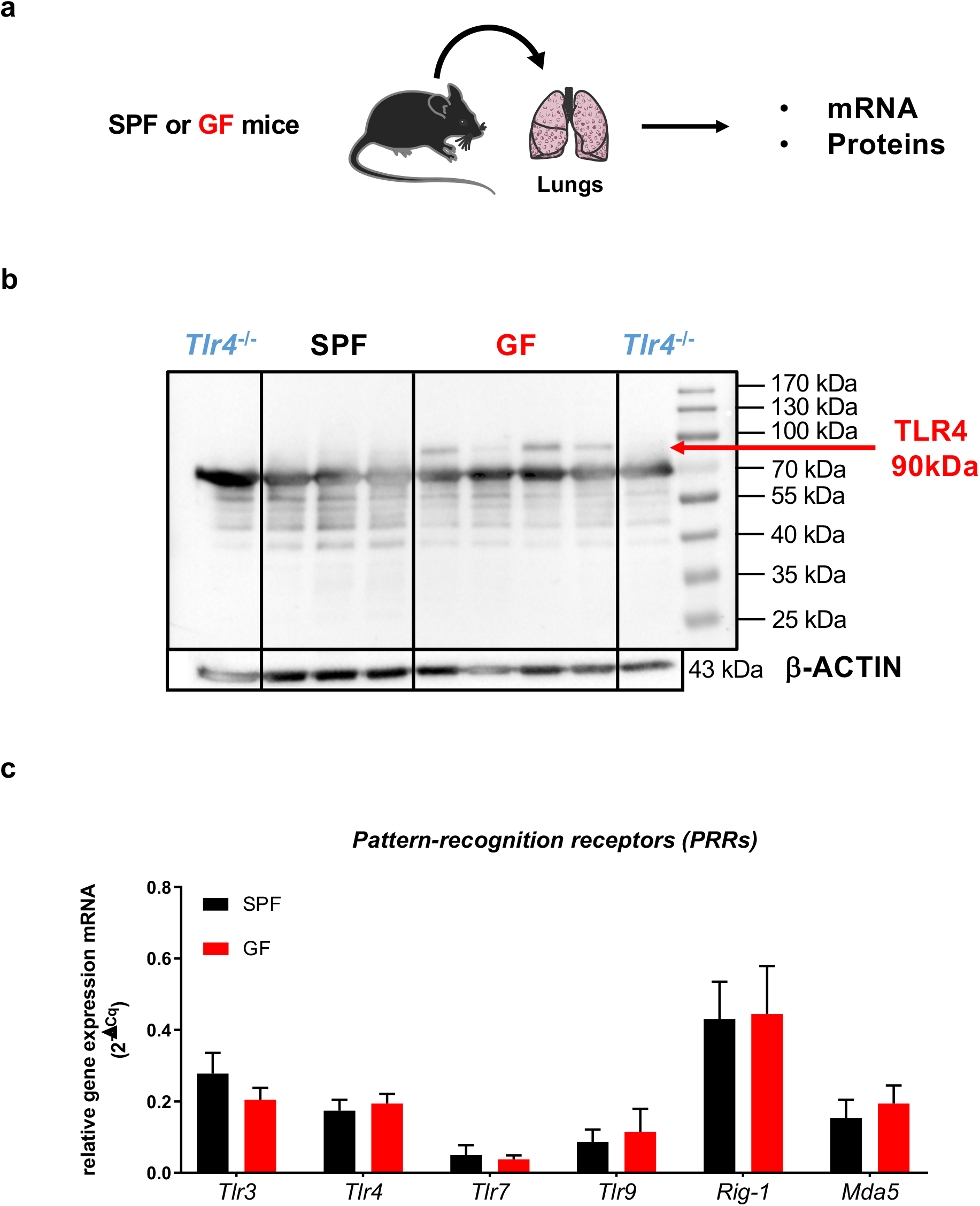
The lungs of GF mice express a higher level of TLR4 at steady state. (**A**) Experimental design. The lungs from GF (n = 4), SPF mice (n = 3) and TLR4-deficient mice (TLR4^-/-^, n = 2) were collected at steady state. (**B**) TLR4 (90 kDa) and β-actin (43 kDa) expression by western blotting. (**C**) The relative mRNA level of various innate sensing receptors was determined by RT-qPCR of lung lysates. Data were analyzed using CFX Manager™ Software (Bio-rad) to determine the quantification cycle (Cq) values. Results were determined using the formula 2^-ΔCq^, with ΔCq = Cqgene-CqHPRT. Results are expressed as the mean ± SD of 6 to 8 biological replicate samples/group and are from two independent experiments (**C**).

### Innate immune reactivity to TLR stimulation is globally higher in GF lung explants

The presence of commensal bacteria may also affect the pulmonary inflammatory reactivity of mice upon sensing *via* several innate immune receptors. We tested this hypothesis by exposing lung explants from GF or SPF mice to a panel of four TLR agonists, namely Poly(I:C) (TLR3 agonist), LPS (TLR4 agonist), Imiquimod (TLR7 agonist), or CpG (TLR9 agonist) (Fig. 3A). Stimulation with these TLR agonists did not induce cytotoxicity in the lung explants after 24h (Fig. S2A), but slight cytotoxicity was observed after 48h (Fig. S2B). TNFα levels were significantly higher in the supernatants of GF explants than SPF explants 24h after stimulation with all TLR-agonists (Fig. 3B). At 48h post-stimulation, only the TNFα response to imiquimod remained higher under GF conditions. There was also strong production of IL-6 by GF explants exposed to all TLR-ligands after 24 and 48h (except LPS after 24h, same level as for SPF) (Fig. 3C). Thus, lung tissues from GF mice showed hyper-production of two major inflammatory cytokines, TNFα and IL-6, in response to four TLR ligands. We next expanded our analysis to cytokines related to other types of innate immune responses (type 1, type 2, type 17/22). The production of IL-12p70, IFNg, TSLP, IL-25, IL-4, IL-5, IL-10, and IL-22 in response to Poly(I:C), LPS, imiquimod, and CpG was quantified in the culture supernatant of lung explants (Fig. S2C and 3D). The GF/SPF ratio was determined for each cytokine and each time point and are presented as a heatmap in which red indicates higher production of the cytokines by GF than SPF explants and blue the inverse (Fig. 3D). Inflammatory and type 1 and type 2 cytokine secretions were notably higher in GF explants 24h following stimulation with Poly(I:C), LPS, and imiquimod, whereas CpG preferentially induced inflammatory and type 1 cytokine release (Fig. 3D and Fig. S2C). Moreover, IL-22 secretion was markedly higher in GF explants at 48h (Fig. 3D and Fig. S2D). Overall, GF lung explants produced more innate immune cytokines in response to a panel of four TLR agonists, thus showing global over-reactivity to pathogen-associated molecular patterns (PAMPs). By contrast, the lungs from mice with a microbiota displayed moderate innate immune responses *ex vivo* when exposed to TLR-dependent stimuli.

**Figure 3.**
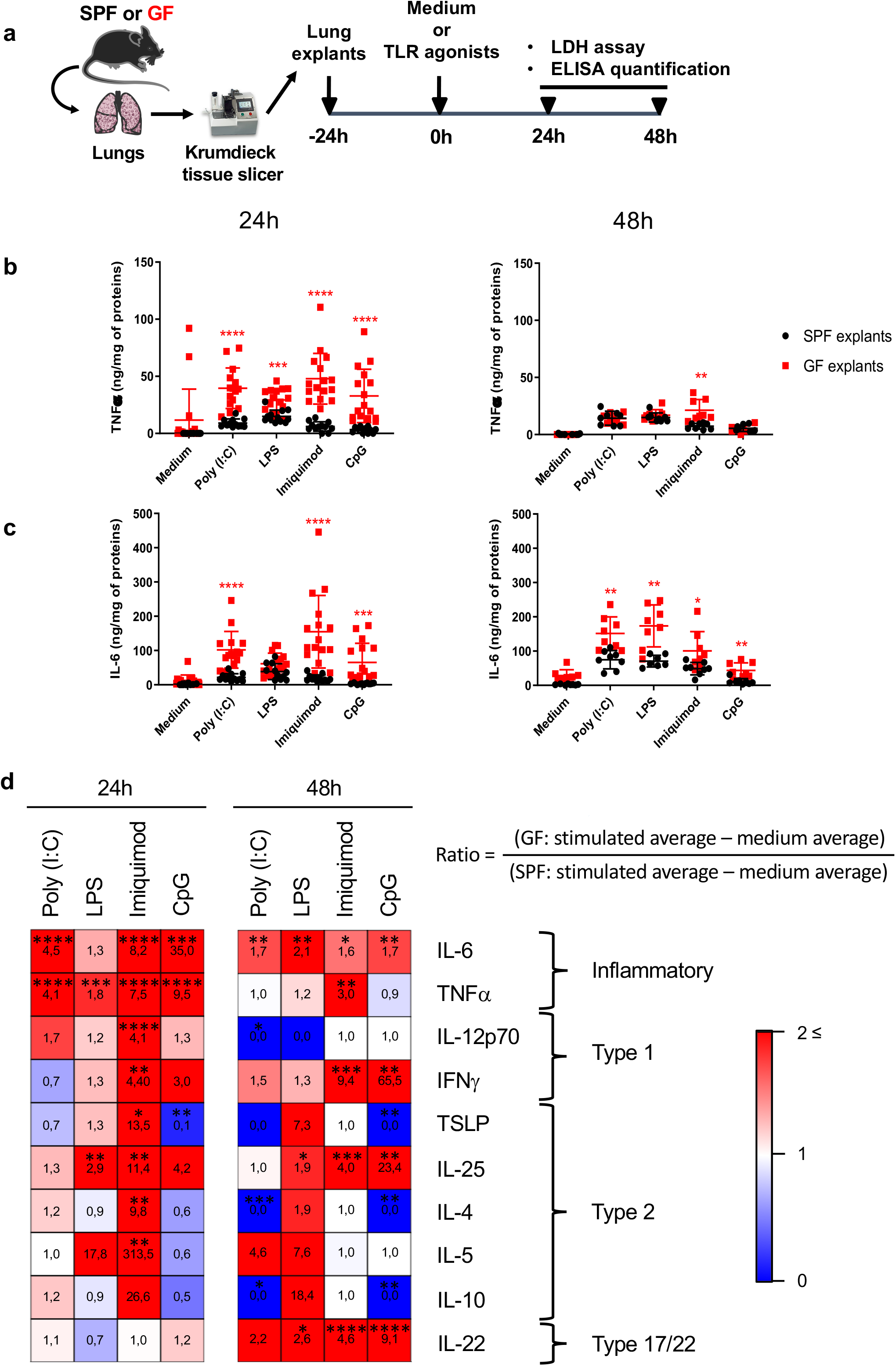
Pulmonary explants from GF mice show increased cytokine production after TLR agonist stimulation. (**A**) Experimental design. The lungs from GF and SPF mice were isolated and cut with a Krumdieck tissue slicer to obtain explants. They were next stimulated with TLR agonists or a medium control for 24 or 48 h and the supernatants collected for cytokine-production analysis. (**B**) TNFα and (**C**) IL-6 production was measured in the supernatants of lung explants. (**D**) Heatmaps showing the ratio (GF/SPF) of cytokine production (IL-12p70, IFNg, TSLP, IL-25, IL-4, IL-5, IL-10, and IL-22) 24 and 48 h after TLR agonist stimulation. Stars determine a significant difference between GF and SPF groups according to histograms in Fig. S2. Results are expressed as the mean ± SD of 4 to 8 biological replicate samples/group and represent a pool of two independent experiments. \**P* ≤ 0.05, \*\**P* ≤ 0.01, \*\*\**P* ≤ 0.001, \*\*\*\**P* ≤ 0.0001 between the SPF and GF groups (Mann-Whitney test) (**B**-**C**).

### Innate immune reactivity is augmented in GF lung explants exposed to RSV

We next extended our observations on the innate immune response of GF mice, obtained with synthetic agonists of TLRs, to a living viral pathogen in order to obtain a physiological relevance of our data. Thus, we investigated the outcome of the GF condition on the pulmonary immune response to a human respiratory virus, such as RSV. GF and SPF lung explants were exposed for 24 or 48h to RSV or a mock control (supernatant of non-infected Hep2 cells used to grow the RSV) (Fig. 4A). Importantly, the cytotoxic effects of the RSV and mock treatments were identically low for the GF and SPF explants after 24h (Fig. S3A). However, after 48h, the cytotoxic impact of RSV and mock treatment was slightly higher in the GF explants (Fig. S3B). TNFα and IL-6 production were higher in the GF explants 24h after exposure to RSV, and to a lesser extent for the mock condition (Fig. 4B). After 48h, TNFα production was lower and the difference between the SPF and GF explants disappeared. IL-6 production was still higher in the GF explants after 48h, both for RSV and the mock control condition (Fig. 4B). The levels of a panel of type 1, type 2, and type 17/22 cytokines were higher (red squares) after 24h in the GF explants (Fig. 4C and Fig. S3C-D) except for IL-25. The differences between GF and SPF were attenuated 48h after exposure to RSV, with an equal or even higher production of cytokines by the SPF explants. These data show that GF lungs mount a stronger and faster cytokine response to RSV exposure than SPF lungs.

**Figure 4.**
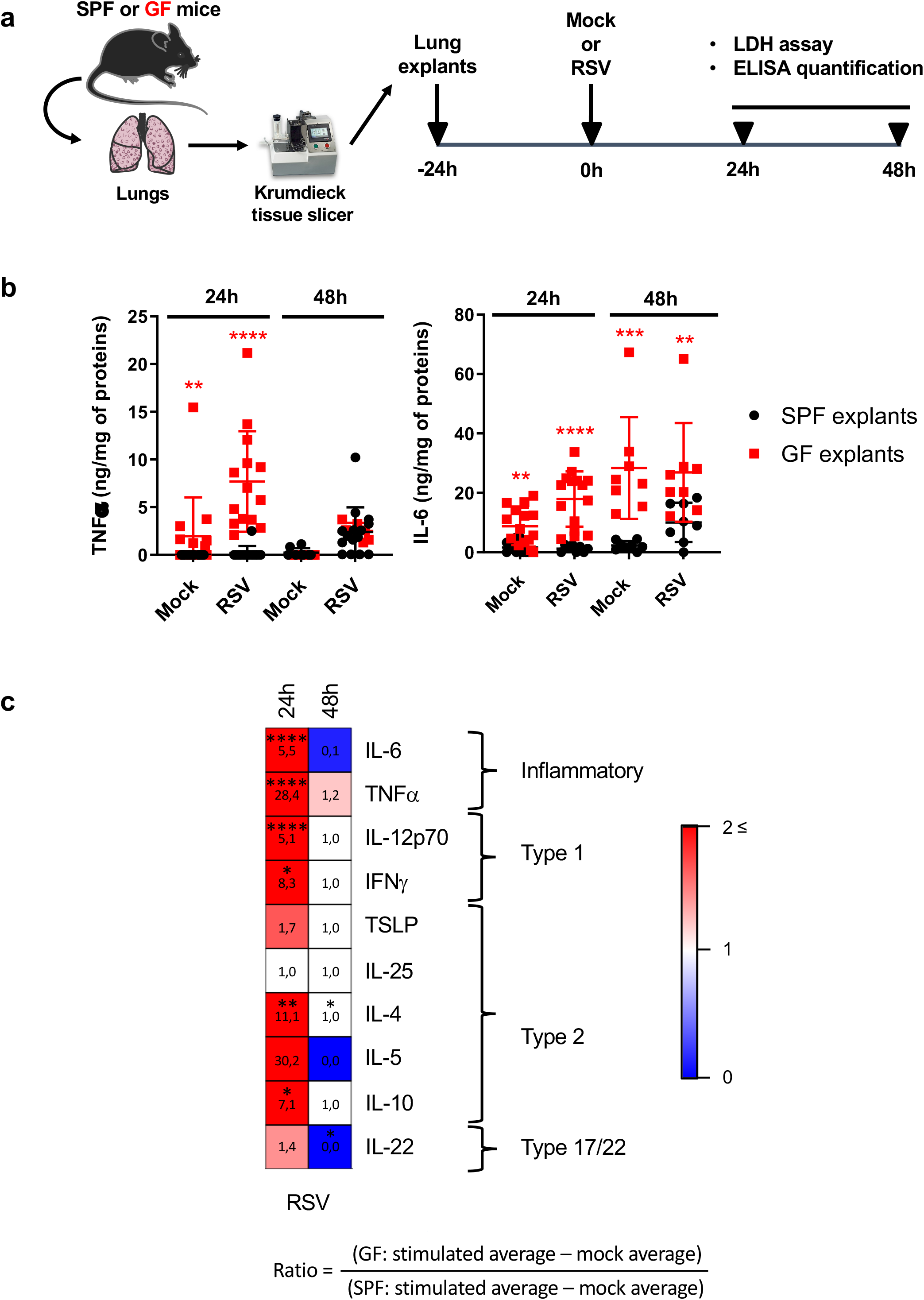
Pulmonary explants from GF mice show increased cytokine production after *ex vivo* RSV infection. (**A**) Experimental design. The lungs from GF and SPF mice were isolated and cut with a Krumdieck tissue slicer to obtain explants. They were next infected with RSV-Cherry or Mock control for 24 or 48 h and the supernatants collected for cytokine-production analysis. (**B**) TNFα and IL-6 production (ng/mg of proteins) was measured in the supernatants of lung explants. (**C**) Heatmap showing the ratio (GF/SPF) of cytokine production (IL-12p70, IFNg, TSLP, IL-25, IL-4, IL-5, IL-10, and IL-22) 24 and 48 h after RSV infection. Stars determine a significant difference between GF and SPF groups according to histograms in Fig. S3. Results are expressed as the mean ± SD of 4 to 8 biological replicate samples/group and represent a pool of two independent experiments. *\*P* ≤ 0.05, \*\**P* ≤ 0.01, \*\*\**P* ≤ 0.001, \*\*\*\**P* ≤ 0.0001 between the SPF and GF groups (Mann-Whitney test) (**B**).

### Alveolar macrophages from GF mice show strong innate immune reactivity, a type-I interferon response, and reduced viral replication

AMs are among the first immune resident cells to encounter RSV and play an important role in the early response to RSV exposure (27). AMs isolated from GF or SPF mice were infected for 24h with RSV (Fig. 5A). GF AMs produced TNFα and KC/CXCL1 in response to RSV infection, whereas SPF AMs did not (Fig. 5B). The mRNA levels of all PRRs tested (*Tlr3, Tlr4, Tlr7, Tlr9, Rig1*, and *Mda5*) were also higher in GF AMs 24h after RSV infection (Fig. 5C). Thus, GF AMs show a high inflammatory profile in response to RSV infection. Interestingly, *Tlr4* mRNA levels were already higher in the Mock-GF control group than the Mock-SPF group (Fig. 5C).

**Figure 5.**
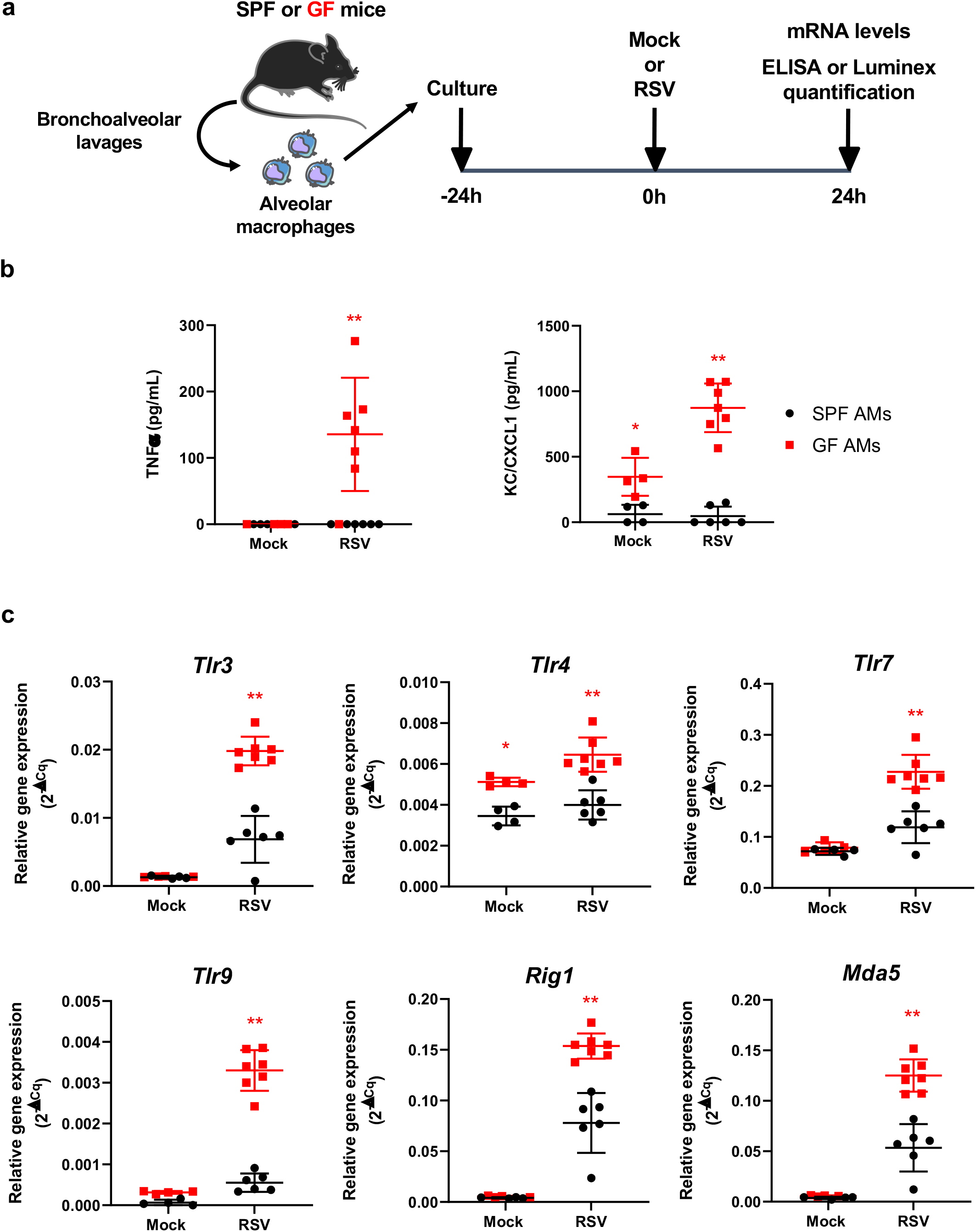
Pro-inflammatory cytokine production and innate sensing receptor expression are higher in RSV-infected alveolar macrophages (AMs) from GF mice. (**A**) Experimental design. AMs were isolated from GF and SPF mice and infected with RSV or control Hep2-supernatant (Mock) at a MOI of 5 for 2 h. (**B**) The production of TNFα and KC/CXCL1 in cell supernatants were measured 24 h post-infection by ELISA. (**C**) The mRNA levels of various innate sensing receptors was determined by RT-qPCR on cell lysates. Results are expressed as the mean ± SD of 4 to 7 biological replicate samples/group and are representative of two independent experiments. \**P* ≤ 0.05 and \*\**P* ≤ 0.01 between the SPF and GF groups (Mann-Whitney Test) (**B**-**C**).

Upon infection with RSV, AMs are the major producer of type-I interferons (IFN-I), which induce the expression of interferon-stimulated genes (ISGs) and activate antiviral pathways in infected cells (28). The production of IFN-I (IFNα and IFNβ) was significantly higher in the supernatants of GF AMs infected with RSV, whereas only IFNβ was produced at detectable levels by RSV-infected SPF AMs (Fig. 6A). The mRNA levels of several ISGs, such as *Ifitm3, Oas, Isg15*, and *Irf7* were also significantly higher in GF AMs than in SPF AMs after RSV infection (Fig. 6B). In accordance with all anti-viral molecules overproduced by GF AMs, the expression of the *N* viral gene, an indicator of RSV replication, was significantly lower in RSV-infected GF AMs than SFP AMs (Fig. 6C). In summary, GF AMs were more efficient at controlling RSV infection than SPF AMs *via* the stronger induction of IFN-I signaling pathways.

**Figure 6.**
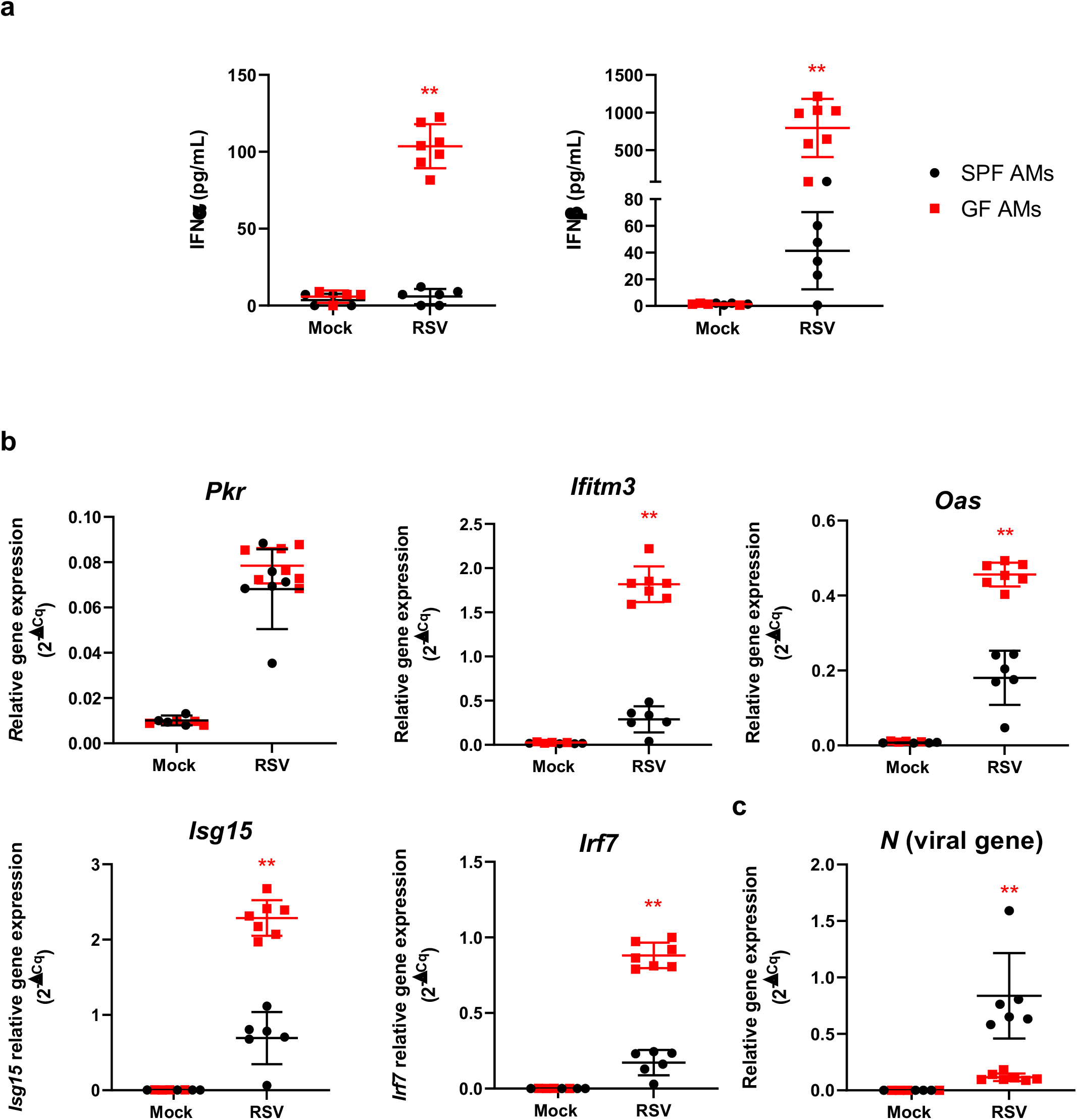
Alveolar macrophages (AMs) from GF mice show increased production of type-I IFNs following *ex vivo* RSV infection. AMs from GF and SPF mice were infected with RSV or control Hep2-supernatant (Mock) at a MOI of 5 for 2 h. (**A**) IFNα and IFNβ production was measured in the supernatants of AMs 24 h after infection. (**B**) *Pkr, Ifitm3, Oas, Isg15, Irf7*, and (**C**) *N* (viral gene) mRNA levels were assessed in cell lysates by RT-qPCR. Data were analyzed using CFX Manager™ Software (Bio-rad) to determine the quantification cycle (Cq) values. Results were determined using the formula 2^-ΔCq^, with ΔCq = Cqgene-CqGADPH. Results are expressed as the mean ± SD of 4 to 7 biological replicate samples/group and are from two independent experiments. \*\**P* ≤ 0.01 between the SPF and GF groups (Mann-Whitney test) (**A**-**C**).

## Discussion

We explored the innate immune function of pulmonary tissue in response to TLR agonist stimulation or RSV infection in a GF *versus* SPF context. The GF lung condition, whether *in vivo* (i.n. LPS administration) or *ex vivo* (lung explants or AMs exposed to TLR ligands or RSV) was consistently associated with elevated expression of inflammatory cytokines and elevated activation of IFN-I pathways. Thus, intranasal instillation of LPS to GF mice resulted in an earlier inflammatory response in the lungs than in SPF mice. This was particularly noticeable for TNFα and KC/CXCL1 production (in BAL and tissue) and neutrophil recruitment in BAL. The IL-6 production kinetic was also accelerated in the BAL of GF mice. Herbst *et al*. also observed a higher inflammatory response, with markers of type-2 immunity, in GF mice when applying a protocol for OVA-induced allergic inflammation (10).

The gut microbiota is known to inhibit TLR signaling pathways through the LPS of the *Bacteroidetes* phylum (29). These findings are consistent with the detectable TLR4 protein in the lungs of GF mice but not (or at undetectable levels by western blot analysis) SPF mice. Thus, the pulmonary innate immune system in mice without bacteria appears to be predisposed to react quickly and strongly to external stimuli. The specificity of commercialized antibodies to detect the expression of other TLRs (*i.e*. TLR3, TLR7, and TLR9) was not sufficient to record potential differences between SPF and GF lungs at steady state (data not shown). Thus, the reactivity of TLR3, TLR7, and TLR9 was assessed functionally in GF lung explants and compared to that of SPF lung explants using their respective agonists. TLR agonist stimulation and RSV infection did not induce tissue mortality as measured by LDH release, except for all GF conditions at 48h. The reactivity of GF explants measured by TNFα and IL-6 production after stimulation with TLR3, TRL4, TLR7, and TLR9 agonists or RSV was much higher than that of SPF explants, notably at 24h. The difference in the kinetics between the IL-6 and TNFα response may be related to a shorter half-life or faster consumption of TNFα by the pulmonary tissue. The background detection of IL-6 was likely related to higher tissue mortality (as measured by LDH release). Analysis of the complete set of cytokines investigated showed a more nuanced response. Indeed, even in the GF context, the production of cytokines varied with the type of stimulus. Imiquimod appeared to be the best inducer of type-1 and type-2 cytokine production, whereas CpG was a poor inducer of type-2 cytokine production in GF lung explants. Most cytokines tested were upregulated at 24h in RSV-infected GF lung explants, but not at 48h. IL-22 production differed strikingly in GF explants depending on the stimulus. All TLR ligands were able to induce high levels of IL-22 at 48h in GF explants, whereas exposure to RSV poorly triggered its production. By contrast, exposure to RSV of GF AMs resulted in an overall hyperactive phenotype relative to that of SPF AMs. Indeed, TLR mRNA levels and those of pro-inflammatory cytokines, IFN-I, and ISGs were upregulated in GF AMs, leading to the better control of RSV infection. Overall, the GF condition showed higher reactivity of the pulmonary innate immune response after stimulation with several TLR agonists and RSV than the SPF condition.

Our data suggest that the presence of a microbiota is associated with a reduction in inflammatory responses, with low TLR4 expression in the lungs of SPF mice and *TLR4* transcripts in SPF AMs. This finding is supported by a study of Didierlaurent *et al*. in which the authors showed that TLR reactivity is diminished for several months after an influenza virus infection. Indeed, secondary stimulation with TLR agonists or bacterial infection leads to a diminution of NF-kB activity and downstream TLR signaling, resulting in reduced inflammatory cytokine production and neutrophil recruitment (30). The reduction of TLR activity induces a mechanism of innate immune tolerance in the lungs, which provides the opportunity for secondary infections to occur, such as with *Pseudomonas aeruginosa* or *Streptococcus pneumonia*. A similar mechanism could explain why more innate immune pathways were activated in the absence of a microbiota (GF lung or GF AM). Without microbiota, TLR expression (notably TLR4) was not repressed, leading to higher reactivity of lung innate immune responses to TLR agonists and RSV infection. Indeed, TLR4 signaling can be activated by the F protein of RSV and act as a co-receptor for viral entry into target cells (31). Recently, TLR4 was reported to play a role in the pathogenesis of RSV disease (32). Given the high expression of TLR4 in GF lungs and high *Tlr4* mRNA levels in AMs, cells may be more permissive to RSV entry and thus become more highly activated than in the SPF condition. Thus, the presence of TLR4 may have contributed to the high reactivity of the pulmonary tissue and AMs to RSV in the GF condition.

Following imprinting with PAMPs, innate immune cells do not behave in the same way to a secondary stimulation or infection (33). This observation refers to an emerging concept called “Innate immune memory”. It involves the reprogramming of innate immune cell metabolism and epigenetic modifications induced by PAMPs that lead to a hyper- or hypo-response of these cells (34). It has been demonstrated that epithelial cells can also undergo epigenetic reprogramming after flagellin stimulation, using histone enzyme inhibitors, and that this conditions their response to *Aspergillus fumigatus* infection (35). In our case, the microbiota, as a priming signal, may have induced this innate immune memory process towards tolerance, explaining the higher reactivity of the pulmonary tissue in its absence (GF condition). Bacterial metabolites from the microbiota, such as short-chain fatty acids (SCFAs), acetate, propionate, and butyrate, may be involved in this innate immune tolerance process. It is known that SCFAs are involved in epigenetic modifications through their inhibitory activity on histone deacetylase (36), making them potentially important for tolerance induction by the microbiota. Moreover, it has been demonstrated that acetate inhibits LPS-induced TNFα production by mononuclear cells from humans and mice through G protein-coupled receptor 43 (GPR43) (37). Butyrate is able to decrease the expression of TLR4 by intestinal epithelial cells and the production of TNFα, monocyte-chemoattractant protein-1 (MCP-1/CCL2), and IL-6 through the activation of GPR41 in RAW264.7 macrophages (38, 39). However, further studies are required to show whether the SCFAs often produced by the gut microbiota are required for innate immune tolerance in the lungs. The question also arises as to which microbiota is involved in the skewing of the pulmonary innate immune system towards tolerance. Deservedly, both the lung and gut microbiota could be involved, as SCFAs systemically circulate from the gut to the lungs through the “gut-lung axis” (40). Imprinting by the microbiota appears to educate the innate immune response of the airways by reducing TLR4 production and this may be true for other TLRs as well. Although microbiota-driven innate immune tolerance may be deleterious upon acute viral respiratory infection, such as by RSV, it would be essential for avoiding chronic inflammation of the tissue and maintaining homeostasis of the airways, in analogy with the gut compartment.

Besides, we still do not know if the pulmonary phenotype observed in this study is a consequence associated locally with the pulmonary microbiota or indirectly with the intestinal microbiota, or both. The ultimate demonstration would be to transfer the microbiota from SPF mice to GF mice to see this causative effect on pulmonary innate immune sensitivity. It would be informative to compare the phenotype obtained after a transfer by gavage of the animals and therefore a primary colonization of the gut, with a transfer by intra-tracheal route for a primary colonization of the lungs. Moreover, based on our previous data obtained in the gut (41), we know that the delay after the colonization is crucial. So, the age of transfer and the delay after transfer require further investigations in particular in the context of the lungs where the colonization and the acquisition of a pulmonary microbiota occur progressively after birth from the oropharyngeal sphere. Those aspects, to our knowledge, have never been investigated and constitute future promising developments.

To conclude, our data suggest the involvement of the microbiota in the tuning of the pulmonary innate immune system. These findings also support the emerging concept of using respiratory commensal bacteria as potential next-generation probiotics to prevent susceptibility to respiratory diseases.

## Abbreviations

AMs: alveolar macrophages
BAL: Bronchoalveolar lavage
COPD: chronic obstructive pulmonary disease
GF: germ-free
GPR43: G-protein-coupled receptor 43
HDM: house dust mites
IFN-I: type 1 interferons
i.n.: intranasal
KC: keratinocyte chemoattractant
LPS: lipopolysaccharide
MCP-1: monocyte chemoattractant protein-1
MGG: May-Grunwald-Giemsa
OVA: ovalbumin
PAMPs: pathogen-associated molecular patterns
pDCs: plasmacytoid dendritic cells
PRRs: patternrecognition receptors
RSV: respiratory syncytial virus
SCFAs: short-chain fatty acids
SPF: specific-pathogen-free

## Acknowledgements

We are grateful to M-A. Rameix-Welti (U1173, UVSQ) and J-F Eléouët (INRAE, UVSQ, VIM) for providing recombinant RSV-Cherry. We are grateful to the IERP unit, INRAE (Infectiology of fishes and rodent facility, doi: 10.15454/1.5572427140471238E12) and ANAXEM in MICALIS unit to animals’ facilities and for birth management. We thank the Emerg’in platform for access to IVIS200 that was financed by the Region Ile De France (SESAME). C. Drajac. and Q. Marquant were the recipient of a Ph.D. fellowship of Région Ile-de-France (DIM-Malinf or DIM-OneHealth, respectively). D. Laubreton was supported by a grant of the French national research agency “Agence Nationale de la Recherche” (ANR-13-BSV3-0016). This project was supported by the Fondation Air Liquide.

## Authors’ contributions

All authors have contributed to, seen, and approved the final, submitted version of the manuscript. Q.M., D.L., S.R., M.T. and D.D. conceptualized the study and designed experiments. Q.M., D.L., C.D., E.M., E.B., M-L.N., A.R. and D.D. performed experiments and generated data analysis. Q.M., D.L., S.R., M.T. and D.D. guided the analysis of these data. Q.M., D.L., S.R., M.T. and D.D. wrote the manuscript with input from all authors.

## Supplemental figure legend

**Figure S1:**
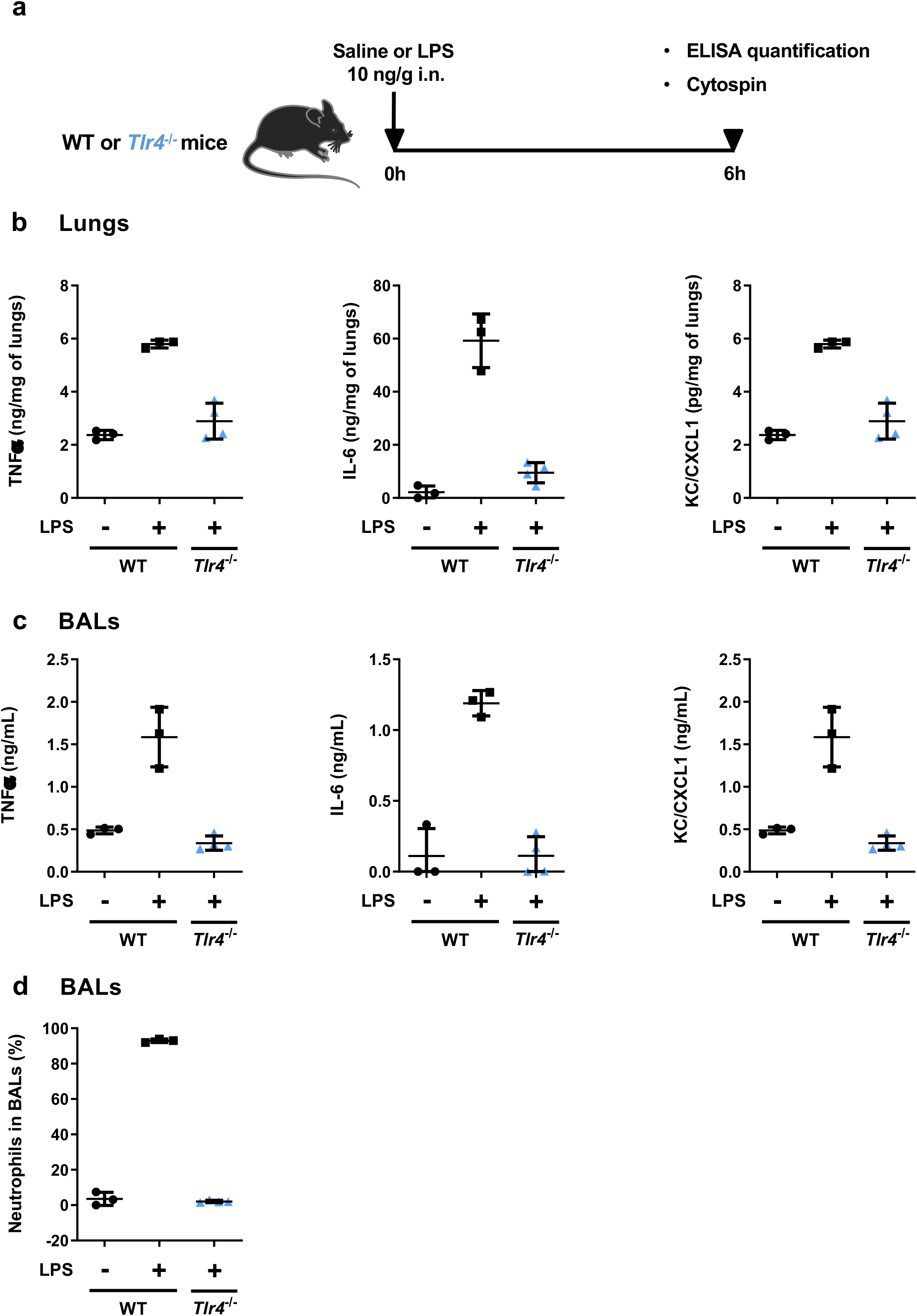
Inflammatory responses in the lungs of wild-type (WT) or TLR4-deficient (TLR4^-/-^) mice after LPS stimulation. (**A**) Experimental design. WT or *Tlr4^-/-^* mice were anesthetized and saline solution or LPS (10 ng/g) was administrated intra-nasally (i.n.). Mice were euthanized 6 h after LPS (10ng/g of mice) administration to evaluate pulmonary pro-inflammatory parameters. (**B-C**) TNFα, IL-6 and KC/CXCL1 production was assessed in BALs (ng/mL) and the lung homogenates (pg/mL or ng/mL) from SPF or TLR4^-/-^ mice. (**D**) The percentage of neutrophils was also determined in BALs. Results are expressed as the mean ± SD (n = 3-4 mice/group).

**Figure S2.**
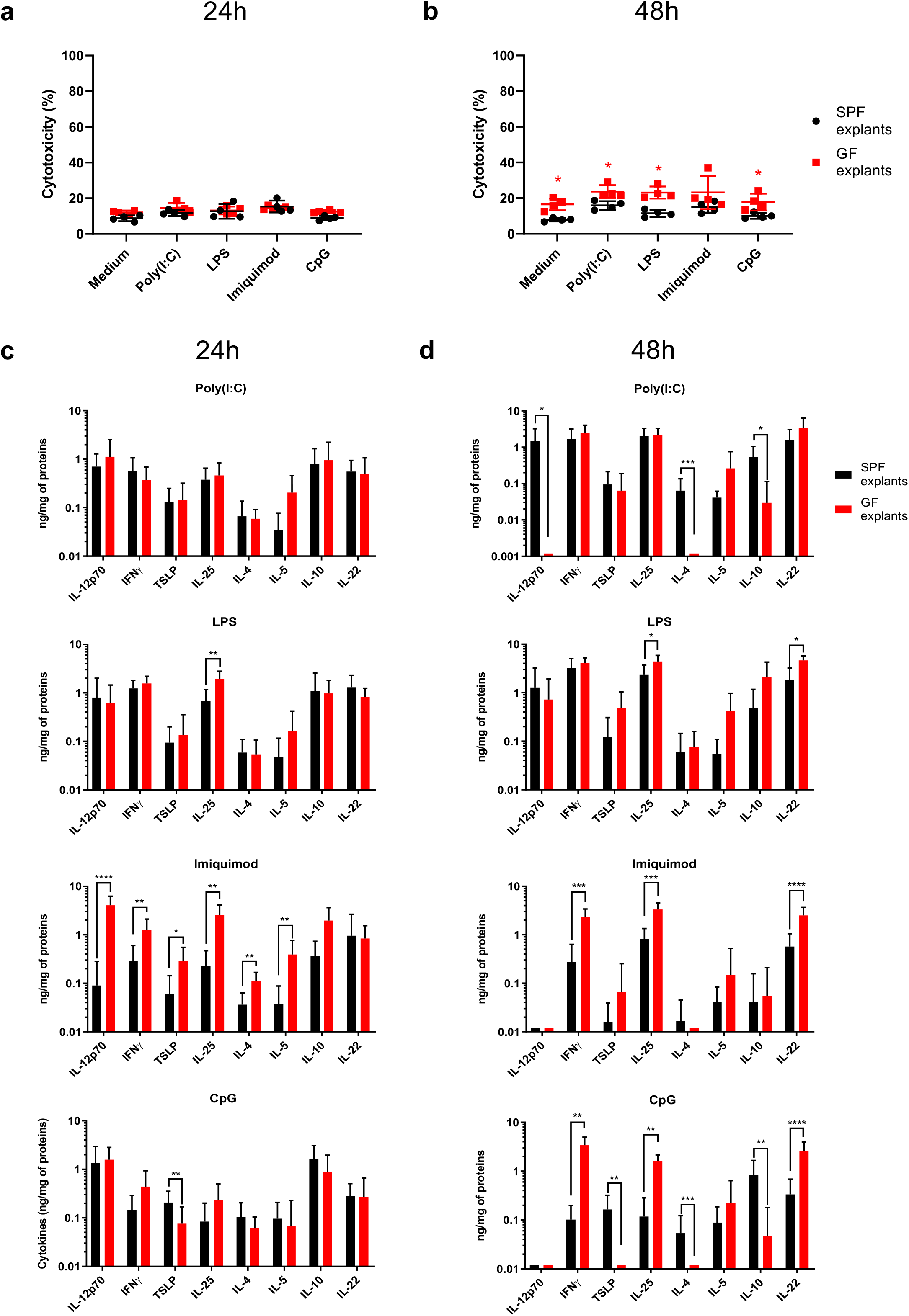
Cytokine production after TLR-agonist stimulation of lung explants from SPF and GF mice. GF and SPF lung explants were obtained and stimulated as described in Figure 3. (**A**) The cytotoxicity in lung explants was measured by the LDH assay 24 h and (**B**) 48 h after stimulation. (**C**) Histograms showing the production of the cytokines IL-12p70, IFNg, TSLP, IL-25, IL-4, IL-5, IL-10, and IL-22 (ng/mg of proteins) 24 h and (**D**) 48 h after TLR-agonist stimulation. The corresponding average medium control value was subtracted from that of each individual sample. Results are expressed as the mean ± SD of 4 to 8 biological replicate samples/group and are from two independent experiments. \**P* ≤ 0.05, \*\**P* ≤ 0.01, \*\*\**P* ≤ 0.001, and \*\*\*\**P* ≤ 0.0001 between the SPF and GF groups (Mann-Whitney test) (**A**-**D**).

**Figure S3.**
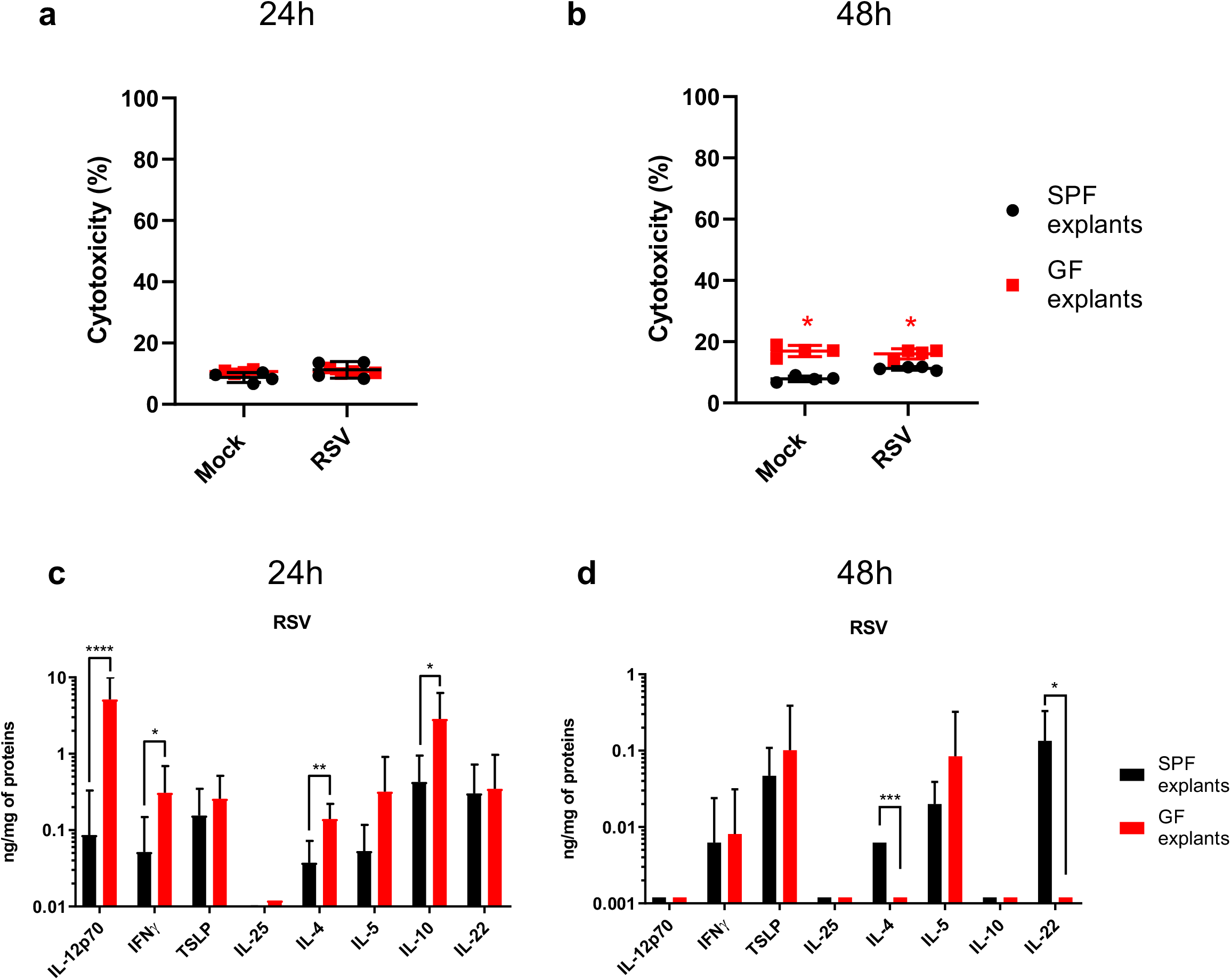
Cytokine production and viral replication after RSV infection of lung explants from SPF and GF mice. GF and SPF lung explants were obtained and stimulated as described in Figure 4. **(A)** The cytotoxicity in lung explants was measured by the LDH assay 24h and **(B)** 48 h after stimulation. **(C)** Histograms showing the production of the cytokines IL-12p70, IFNg, TSLP, IL-25, IL-4, IL-5, IL-10, and IL-22 (ng/mg of proteins) 24 h and **(D)** 48 h after RSV infection. The corresponding average medium control value was subtracted from that of each individual sample. Results are expressed as the mean ± SD of 4 to 8 biological replicates samples/group and are from two independent experiments. \**P* ≤ 0.05, \*\**P* ≤ 0.01, \*\*\**P* ≤ 0.001, and \*\*\*\**P* ≤ 0.0001 between the SPF and GF groups (Mann-Whitney test) **(A-D)**.

## References

1. van de Guchte, M., Blottiere, H. M., and Dore, J. (2018) Humans as holobionts: implications for prevention and therapy. Microbiome 6, 81

2. Takiishi, T., Fenero, C. I. M., and Camara, N. O. S. (2017) Intestinal barrier and gut microbiota: Shaping our immune responses throughout life. Tissue Barriers 5, e1373208

3. Sommer, F., and Backhed, F. (2013) The gut microbiota--masters of host development and physiology. Nat Rev Microbiol 11, 227–238

4. Biesbroek, G., Tsivtsivadze, E., Sanders, E. A., Montijn, R., Veenhoven, R. H., Keijser, B. J., and Bogaert, D. (2014) Early respiratory microbiota composition determines bacterial succession patterns and respiratory health in children. Am J Respir Crit Care Med 190, 1283–1292

5. Remot, A., Descamps, D., Noordine, M. L., Boukadiri, A., Mathieu, E., Robert, V., Riffault, S., Lambrecht, B., Langella, P., Hammad, H., and Thomas, M. (2017) Bacteria isolated from lung modulate asthma susceptibility in mice. ISME J 11, 1061–1074

6. Huang, Y. J., Charlson, E. S., Collman, R. G., Colombini-Hatch, S., Martinez, F. D., and Senior, R. M. (2013) The role of the lung microbiome in health and disease. A National Heart, Lung, and Blood Institute workshop report. Am J Respir Crit Care Med 187, 1382–1387

7. Venkataraman, A., Bassis, C. M., Beck, J. M., Young, V. B., Curtis, J. L., Huffnagle, G. B., and Schmidt, T. M. (2015) Application of a neutral community model to assess structuring of the human lung microbiome. MBio 6

8. Dickson, R. P., Erb-Downward, J. R., Freeman, C. M., McCloskey, L., Falkowski, N. R., Huffnagle, G. B., and Curtis, J. L. (2017) Bacterial Topography of the Healthy Human Lower Respiratory Tract. mBio 8

9. Pulvirenti, G., Parisi, G. F., Giallongo, A., Papale, M., Manti, S., Savasta, S., Licari, A., Marseglia, G. L., and Leonardi, S. (2019) Lower Airway Microbiota. Front Pediatr 7, 393

10. Herbst, T., Sichelstiel, A., Schar, C., Yadava, K., Burki, K., Cahenzli, J., McCoy, K., Marsland, B. J., and Harris, N. L. (2011) Dysregulation of allergic airway inflammation in the absence of microbial colonization. Am J Respir Crit Care Med 184, 198–205

11. Huang, Y. J., Nariya, S., Harris, J. M., Lynch, S. V., Choy, D. F., Arron, J. R., and Boushey, H. (2015) The airway microbiome in patients with severe asthma: Associations with disease features and severity. J Allergy Clin Immunol 136, 874–884

12. Marri, P. R., Stern, D. A., Wright, A. L., Billheimer, D., and Martinez, F. D. (2013) Asthma-associated differences in microbial composition of induced sputum. J Allergy Clin Immunol 131, 346–352 e341-343

13. Jain, S., Williams, D. J., Arnold, S. R., Ampofo, K., Bramley, A. M., Reed, C., Stockmann, C., Anderson, E. J., Grijalva, C. G., Self, W. H., Zhu, Y., Patel, A., Hymas, W., Chappell, J. D., Kaufman, R. A., Kan, J. H., Dansie, D., Lenny, N., Hillyard, D. R., Haynes, L. M., Levine, M., Lindstrom, S., Winchell, J. M., Katz, J. M., Erdman, D., Schneider, E., Hicks, L. A., Wunderink, R. G., Edwards, K. M., Pavia, A. T., McCullers, J. A., Finelli, L., and Team, C. E. S. (2015) Community-acquired pneumonia requiring hospitalization among U.S. children. N Engl J Med 372, 835–845

14. de Steenhuijsen Piters, W. A., Heinonen, S., Hasrat, R., Bunsow, E., Smith, B., Suarez-Arrabal, M. C., Chaussabel, D., Cohen, D. M., Sanders, E. A., Ramilo, O., Bogaert, D., and Mejias, A. (2016) Nasopharyngeal Microbiota, Host Transcriptome, and Disease Severity in Children with Respiratory Syncytial Virus Infection. Am J Respir Crit Care Med 194, 1104–1115

15. Ege, M. J., Mayer, M., Normand, A. C., Genuneit, J., Cookson, W. O., Braun-Fahrlander, C., Heederik, D., Piarroux, R., von Mutius, E., and Group, G. T. S. (2011) Exposure to environmental microorganisms and childhood asthma. N Engl J Med 364, 701–709

16. Stein, M. M., Hrusch, C. L., Gozdz, J., Igartua, C., Pivniouk, V., Murray, S. E., Ledford, J. G., Marques Dos Santos, M., Anderson, R. L., Metwali, N., Neilson, J. W., Maier, R. M., Gilbert, J. A., Holbreich, M., Thorne, P. S., Martinez, F. D., von Mutius, E., Vercelli, D., Ober, C., and Sperling, A. I. (2016) Innate Immunity and Asthma Risk in Amish and Hutterite Farm Children. N Engl J Med 375, 411–421

17. Caballero, M. T., Serra, M. E., Acosta, P. L., Marzec, J., Gibbons, L., Salim, M., Rodriguez, A., Reynaldi, A., Garcia, A., Bado, D., Buchholz, U. J., Hijano, D. R., Coviello, S., Newcomb, D., Bellabarba, M., Ferolla, F. M., Libster, R., Berenstein, A., Siniawaski, S., Blumetti, V., Echavarria, M., Pinto, L., Lawrence, A., Ossorio, M. F., Grosman, A., Mateu, C. G., Bayle, C., Dericco, A., Pellegrini, M., Igarza, I., Repetto, H. A., Grimaldi, L. A., Gudapati, P., Polack, N. R., Althabe, F., Shi, M., Ferrero, F., Bergel, E., Stein, R. T., Peebles, R. S., Boothby, M., Kleeberger, S. R., and Polack, F. P. (2015) TLR4 genotype and environmental LPS mediate RSV bronchiolitis through Th2 polarization. J Clin Invest 125, 571–582

18. Lloyd, C. M., and Marsland, B. J. (2017) Lung Homeostasis: Influence of Age, Microbes, and the Immune System. Immunity 46, 549–561

19. Kanmani, P., Clua, P., Vizoso-Pinto, M. G., Rodriguez, C., Alvarez, S., Melnikov, V., Takahashi, H., Kitazawa, H., and Villena, J. (2017) Respiratory Commensal Bacteria Corynebacterium pseudodiphtheriticum Improves Resistance of Infant Mice to Respiratory Syncytial Virus and Streptococcus pneumoniae Superinfection. Front Microbiol 8, 1613

20. Ortiz Moyano, R., Raya Tonetti, F., Tomokiyo, M., Kanmani, P., Vizoso-Pinto, M. G., Kim, H., Quilodran-Vega, S., Melnikov, V., Alvarez, S., Takahashi, H., Kurata, S., Kitazawa, H., and Villena, J. (2020) The Ability of Respiratory Commensal Bacteria to Beneficially Modulate the Lung Innate Immune Response Is a Strain Dependent Characteristic. Microorganisms 8

21. Sencio, V., Barthelemy, A., Tavares, L. P., Machado, M. G., Soulard, D., Cuinat, C., Queiroz-Junior, C. M., Noordine, M. L., Salome-Desnoulez, S., Deryuter, L., Foligne, B., Wahl, C., Frisch, B., Vieira, A. T., Paget, C., Milligan, G., Ulven, T., Wolowczuk, I., Faveeuw, C., Le Goffic, R., Thomas, M., Ferreira, S., Teixeira, M. M., and Trottein, F. (2020) Gut Dysbiosis during Influenza Contributes to Pulmonary Pneumococcal Superinfection through Altered Short-Chain Fatty Acid Production. Cell Rep 30, 2934–2947 e2936

22. Gollwitzer, E. S., Saglani, S., Trompette, A., Yadava, K., Sherburn, R., McCoy, K. D., Nicod, L. P., Lloyd, C. M., and Marsland, B. J. (2014) Lung microbiota promotes tolerance to allergens in neonates via PD-L1. Nat Med 20, 642–647

23. Mathieu, E., Escribano-Vazquez, U., Descamps, D., Cherbuy, C., Langella, P., Riffault, S., Remot, A., and Thomas, M. (2018) Paradigms of Lung Microbiota Functions in Health and Disease, Particularly, in Asthma. Front Physiol 9, 1168

24. Rameix-Welti, M. A., Le Goffic, R., Herve, P. L., Sourimant, J., Remot, A., Riffault, S., Yu, Q., Galloux, M., Gault, E., and Eleouet, J. F. (2014) Visualizing the replication of respiratory syncytial virus in cells and in living mice. Nat Commun 5, 5104

25. Descamps, D., Le Gars, M., Balloy, V., Barbier, D., Maschalidi, S., Tohme, M., Chignard, M., Ramphal, R., Manoury, B., and Sallenave, J. M. (2012) Toll-like receptor 5 (TLR5), IL-1beta secretion, and asparagine endopeptidase are critical factors for alveolar macrophage phagocytosis and bacterial killing. Proc Natl Acad Sci U S A 109, 1619–1624

26. Xiang, M., and Fan, J. (2010) Pattern recognition receptor-dependent mechanisms of acute lung injury. Mol Med 16, 69–82

27. Makris, S., Bajorek, M., Culley, F. J., Goritzka, M., and Johansson, C. (2016) Alveolar Macrophages Can Control Respiratory Syncytial Virus Infection in the Absence of Type I Interferons. J Innate Immun 8, 452–463

28. Goritzka, M., Makris, S., Kausar, F., Durant, L. R., Pereira, C., Kumagai, Y., Culley, F. J., Mack, M., Akira, S., and Johansson, C. (2015) Alveolar macrophage-derived type I interferons orchestrate innate immunity to RSV through recruitment of antiviral monocytes. J Exp Med 212, 699–714

29. d’Hennezel, E., Abubucker, S., Murphy, L. O., and Cullen, T. W. (2017) Total Lipopolysaccharide from the Human Gut Microbiome Silences Toll-Like Receptor Signaling. mSystems 2

30. Didierlaurent, A., Goulding, J., Patel, S., Snelgrove, R., Low, L., Bebien, M., Lawrence, T., van Rijt, L. S., Lambrecht, B. N., Sirard, J. C., and Hussell, T. (2008) Sustained desensitization to bacterial Toll-like receptor ligands after resolution of respiratory influenza infection. J Exp Med 205, 323–329

31. Kurt-Jones, E. A., Popova, L., Kwinn, L., Haynes, L. M., Jones, L. P., Tripp, R. A., Walsh, E. E., Freeman, M. W., Golenbock, D. T., Anderson, L. J., and Finberg, R. W. (2000) Pattern recognition receptors TLR4 and CD14 mediate response to respiratory syncytial virus. Nat Immunol 1, 398–401

32. Marzec, J., Cho, H. Y., High, M., McCaw, Z. R., Polack, F., and Kleeberger, S. R. (2019) Toll-like receptor 4-mediated respiratory syncytial virus disease and lung transcriptomics in differentially susceptible inbred mouse strains. Physiol Genomics 51, 630–643

33. Hamon, M. A., and Quintin, J. (2016) Innate immune memory in mammals. Semin Immunol 28, 351–358

34. Boraschi, D., and Italiani, P. (2018) Innate Immune Memory: Time for Adopting a Correct Terminology. Front Immunol 9, 799

35. Bigot, J., Guillot, L., Guitard, J., Ruffin, M., Corvol, H., Chignard, M., Hennequin, C., and Balloy, V. (2020) Respiratory Epithelial Cells Can Remember Infection: A Proof-of-Concept Study. J Infect Dis 221, 1000–1005

36. Correa-Oliveira, R., Fachi, J. L., Vieira, A., Sato, F. T., and Vinolo, M. A. (2016) Regulation of immune cell function by short-chain fatty acids. Clin Transl Immunology 5, e73

37. Masui, R., Sasaki, M., Funaki, Y., Ogasawara, N., Mizuno, M., Iida, A., Izawa, S., Kondo, Y., Ito, Y., Tamura, Y., Yanamoto, K., Noda, H., Tanabe, A., Okaniwa, N., Yamaguchi, Y., Iwamoto, T., and Kasugai, K. (2013) G protein-coupled receptor 43 moderates gut inflammation through cytokine regulation from mononuclear cells. Inflamm Bowel Dis 19, 2848–2856

38. Lee, S. K., Il Kim, T., Kim, Y. K., Choi, C. H., Yang, K. M., Chae, B., and Kim, W. H. (2005) Cellular differentiation-induced attenuation of LPS response in HT-29 cells is related to the down-regulation of TLR4 expression. Biochem Biophys Res Commun 337, 457–463

39. Ohira, H., Fujioka, Y., Katagiri, C., Mamoto, R., Aoyama-Ishikawa, M., Amako, K., Izumi, Y., Nishiumi, S., Yoshida, M., Usami, M., and Ikeda, M. (2013) Butyrate attenuates inflammation and lipolysis generated by the interaction of adipocytes and macrophages. J Atheroscler Thromb 20, 425–442

40. Dang, A. T., and Marsland, B. J. (2019) Microbes, metabolites, and the gut-lung axis. Mucosal Immunol 12, 843–850

41. Tomas, J., Wrzosek, L., Bouznad, N., Bouet, S., Mayeur, C., Noordine, M. L., Honvo-Houeto, E., Langella, P., Thomas, M., and Cherbuy, C. (2013) Primocolonization is associated with colonic epithelial maturation during conventionalization. FASEB J 27, 645–655

